# Variable processing shifts during perceptual acceleration: Evidence from temporal integration

**DOI:** 10.1101/2025.02.12.637809

**Authors:** Michael J. Wolff, Elkan G. Akyürek

## Abstract

The perception of a stimulus can be accelerated by another that precedes it. Perceptual acceleration has been observed in a range of tasks, at varying timescales, and arises by virtue of providing advance spatial and/or temporal information about upcoming stimuli. Here we examined perceptual acceleration during visual temporal integration. Temporal integration occurs when successive stimuli appear that fit together in time as well as space. As such, stimuli arriving first during temporal integration partially predict those that follow. Although temporal integration is a rapid process, we reasoned that this information may cause perceptual acceleration during temporal integration. We used multivariate pattern analysis of EEG data from a missing element task, designed to measure the visual temporal integration of two successive stimulus displays, so that we were able to precisely track the representation associated with the integrated percept in time. We manipulated the delay between our displays, and observed commensurate acceleration of the resultant integrated representation. The degree of acceleration first increased from early (100 ms after stimulus onset) to intermediate (200 ms) processing stages, before decreasing again at a later stage (400 ms). The results thus suggest that perceptual acceleration occurs during temporal integration, but is nonlinear, such that some time that is gained at one moment in the process can be lost again at another.

## Introduction

We live in the past. As we experience the world around us, our impressions inevitably lag behind the physical reality, because it takes some time to route a signal from our eyes and through the brain. For instance, it takes approximately 120-150 ms to determine basic properties of an unfamiliar natural scene, such as to determine whether it contains an animal or not (Cichy et al., 2014; Kirchner & Thorpe, 2006; Thorpe et al., 1996; VanRullen & Thorpe, 2001). Before that delay has elapsed, higher-level awareness of the contents of the scene is considered to be virtually absent, precluding an appropriate behavioural reaction until perceptual processing is complete. This processing time is nevertheless variable, and can be reduced by decreasing the complexity of the scene and the associated task. Furthermore, processing may also be accelerated, if circumstances allow for increased readiness.

Evidence from a range of experimental paradigms has shown perceptual acceleration, typically as a result of providing observers with information about the where and when of upcoming stimuli to prepare for their arrival. Examples of the former include the Posner cueing task (Posner, 1980), in which the reaction time to a target whose location was validly cued beforehand is reduced, and so-called rapid resumption in visual search, where search is faster in displays that were previewed before (Lleras et al., 2005; Spaak et al., 2016). Examples of the latter are studies using temporal cues (Coull & Nobre, 1998), rhythmic presentation sequences (Jones et al., 2002; Mathewson et al., 2010), and predictable foreperiods (Luce, 1991), all of which similarly reduce target reaction time. Here we hypothesized that such perceptual acceleration may also occur during temporal integration.

Temporal integration is thought to take place during perception, in particular when rapid, successive stimuli fit together, such that they can be combined into a single coherent percept (for a review, see Akyürek, 2025). This is exemplified in the missing element task (MET), which is used to measure visual temporal integration (Akyürek et al., 2010; Di Lollo, 1977, 1980), and which we also used here. The MET commonly entails the presentation of two successive displays, each containing a number of stimuli, which are typically simple shapes such as small squares, circles, or dots, laid out in a regularly spaced grid. Each display only shows half of the stimuli, so that together, the two displays fill up the full grid, apart from a single location in the grid that remains empty (see Figure 1). If this location is found by the observer, this is evidence that the successive stimulus displays were temporally integrated, as working out this location from a mental comparison of the individual displays is very unlikely to succeed.

**Figure 1.**
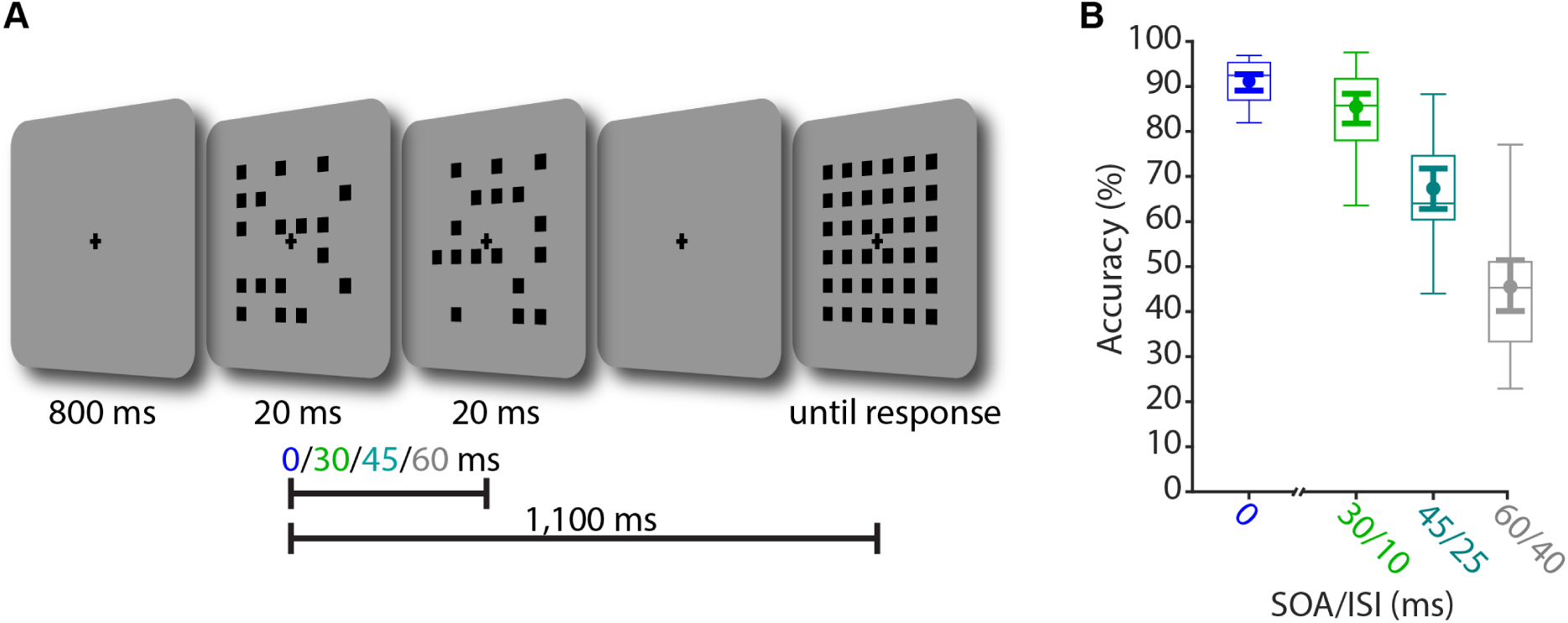
Task structure and behavioral performance (A) Trial schematic of the missing element task (MET). After the presentation of an initial fixation cross, 35 out of 36 squares on a 6 by 6 grid were presented either at the same time (SOA 0 ms) or across two successive displays (SOA 30, 45, and 60 ms). The response screen was presented after a short delay and participants clicked on the square they thought was previously missing. (B) Boxplots show task accuracy as a function of SOA condition. Centre lines indicate the median; box outlines show 25th and 75th percentiles, and whiskers indicate 1.5x the interquartile range. The superimposed filled circles and error bars indicate means and the 95% C.I. of the means, respectively.

Although the stimulus displays in the MET are only very briefly shown, within less than 200 ms in total, the first display does provide both spatial and temporal information as to the appearance of the second display. The locations left empty in the first display will be largely filled by the second display (minus the missing element location), and the first display also signals that the second display is going to appear right after. While stimulus timing is very different in the spatial and temporal cueing tasks mentioned above, in those tasks such advance information is known to accelerate perceptual processing. It is thus conceivable that this might also occur in the MET, more so as rapid interactions between its stimuli have some neural plausibility. For instance, one might speculate that the rapid succession of the mutually compatible stimulus displays may cause coincidental neuronal activity, leading to synaptic facilitation through residual elevation of presynaptic Ca^2+^ (Markram et al., 1997; Zucker & Regehr, 2002). Alternatively, and again speculatively, processing of the first stimulus display in higher-order cells may facilitate processing of the second display in lower-order cells, through back-projections, some of which could operate at MET timescales already (Friston, 2005; Palmer et al., 2015; Summerfield & Egner, 2009).

Using EEG, we could not directly arbitrate between these (or other) neural mechanisms. However, we were able to chart the time course of perceptual acceleration by using multivariate pattern analysis (MVPA) to track the representation of the integrated percept in the brain (Marti et al., 2015; van Ede et al., 2018), as a first step in helping to constrain the range of possible underlying neural mechanisms. Other than expecting evidence for a degree of perceptual acceleration during temporal integration, we did not have strong expectations as to its precise time course. Evidence from cueing, rhythm, and foreperiod studies does not strongly suggest a particular temporal locus. Spatial cueing affects early perceptual processing stages, as it is accompanied by amplitude (but not latency) modulations of the P1 and N1 components of the ERP (Mangun, 1995; Mangun & Hillyard, 1991). Advance temporal information has been found to modulate the N1 (Seibold & Rolke, 2014), N2 (Seibold, Fiedler, et al., 2011), P3 (Griffin et al., 2002), as well as slow potentials and oscillatory components (Praamstra et al., 2006; Rohenkohl & Nobre, 2011). Apart from this already wide temporal range, it is difficult to relate univariate electrophysiological measures to perceptual acceleration directly, as they reflect both perceptual and cognitive processing as well as the representations they act on. Thus, we did not constrain our analyses to a specific time-window, but relied on MVPA to focus on the representation of the missing element itself, to chart when that representation arises, as a function of our task parameters.

### The current study

As indicated, we used a MET to assess perceptual acceleration. We manipulated the stimulus onset asynchrony (SOA) between the displays in our MET, measured the EEG, applied multivariate pattern analysis (MVPA) to decode the location of the missing element, and compared its occurrence in time between conditions by means of cross-temporal generalization. To preview the results, we found evidence for accelerated perceptual processing during temporal integration, but also observed that this acceleration varied over time as the missing element was being processed in the brain, such that initial gains were partially lost again later in that process.

## Materials and Methods

### Participants

Twenty-four first year psychology students of the University of Groningen (9 female, mean age 21 years, range 18-28 years) were included in the analyses. Eight additional participants failed to meet the previously set inclusion criterion (80 % accuracy on the SOA 0/no integration condition), and their data was discarded during collection. The final sample- size of 24 was decided on before data collection and is based on a previous study that used a similar EEG analysis pipeline to decode visual stimuli (Wolff et al., 2015). Participants received course credits for participation and gave written informed consent. The study was approved by the ethical committee of the Psychology department of the University of Groningen (approval number: 16340-S-NE).

### Apparatus and Stimuli

Stimuli were controlled and generated with Psychtoolbox, a freely available toolbox for MATLAB (Kleiner et al., 2007), and presented on a 19-inch (48.3 cm) CRT screen running at 200 Hz refresh rate and a resolution of 640 by 480 pixels. Responses were made with a conventional computer mouse. Participants were seated at approximately 64 cm from the monitor. The stimuli consisted of black squares (0.72°) arranged in a 6 by 6 grid (7.92°) in the center of the screen. The empty spaces between squares were the same size as the squares themselves (0.72°). A black fixation cross (0.97°) and a grey background (RGB = 150, 150, 150) were maintained throughout the trials.

### Procedure

Each trial began with the presentation of a fixation cross. After 800 ms, 35 out of 36 black squares, arranged on a 6 by 6 grid, were presented either at the same time (SOA 0 ms), or across two successive displays, for 20 ms each, with an interstimulus interval (ISI) of 10, 25, or 40 ms (SOA 30, 45, or 60 ms). Participants were instructed to locate the one missing square, which could be anywhere within the inner 4 by 4 grid, and of which they were made aware beforehand. Note that in order to detect the missing square, both displays need to be temporally integrated by the perceptual system, which is more difficult at longer SOAs (Di Lollo, 1977; Hogben & Lollo, 1974). The response screen was presented 1,100 ms after the onset of the first display, and consisted of the whole grid of squares. Participants used the mouse cursor to click on the square that they thought was previously missing. Immediately after their response, the correct square flashed in green, and, if incorrect, the chosen square in red. Participants completed 1,056 trials in total, over a course of approximately 75 minutes, including breaks. See Figure 1a for a trial schematic.

No integration (SOA 0) trials made up 27%, SOA 30 trials made up 27 %, and SOA 45 and SOA 60 made up 36% and 9% of all trials, respectively. SOA 45 made up the largest proportion of trials due to the anticipated difficulty of this condition. Of the most difficult condition of SOA 60, only a small number of trials was included in the experiment. These served merely to confirm the expected decrease in accuracy as a function of SOA due to temporal integration eventually breaking down, as has previously been observed (e.g. Akyürek et al., 2010), and this SOA was excluded from all neurophysiological analyses.

### Electrophysiological recording and pre-processing

The EEG signal was acquired from 59 Ag/AgCls sintered electrodes laid out according to the extended international 10-20 system (Supplemental Figure 1A), with two additional electrodes placed on the mastoids. Another electrode, placed above the sternum, was used as ground. Four electrodes placed above and below the left eye and on the temples recorded the bipolar EOG signals. The data were recorded with a TMSI Refa8-64/72 amplifier, sampling at 1024 Hz with an average reference. Impedances were kept below 10 kΩ. Offline, the data were re-referenced to the average of both mastoid electrodes and filtered with a 0.05 Hz high-pass and a 40 Hz low-pass filter. Only the 24 posterior EEG channels (P7, P5, P3, P1, Pz, P2, P4, P6, P8, PO7, PO3, POz, PO4, PO8, O1, Oz, O2, P9, PO9, O9, Iz, O10, PO10, and P10) were included in all following pre-processing steps and analyses (unless otherwise specified), as we expected the relevant visual signal to be the strongest in these channels, which cover parietal and occipital cortex. This decision was made a-priori and based on our previous experience involving visual tasks and multivariate pattern analyses, where also only the posterior channels were included (e.g., Wolff et al., 2015, 2017, 2020), which is a practice also used by others (e.g., Harrison et al., 2023; Kandemir & Olivers, 2024). The data were epoched relative to the onset of the first display (-250 ms to 1100 ms) and baseline corrected using the mean signal from -250 to 0 ms relative to Display 1 onset. Noisy trials and trials containing ocular artefacts were identified by visually inspecting the EEG and EOG signal and removed from the analyses.

### Location decoding

We were interested in the time-course of the location code of the missing square in the EEG signal. We used Mahalanobis distance (De Maesschalck et al., 2000) to test if the signal contained information about the location at any one point throughout the trial. We used custom MATLAB code as well as functions from the “Statistics and Machine Learning Toolbox” for MATLAB. The above-listed 24 posterior channels were included in the decoding analyses. In essence, we tested if same-location trials were closer to each other in multidimensional space (i.e., a shorter Mahalanobis distance) than different-location trials, where each EEG channel is a separate dimension. Mahalanobis distance is similar to Euclidean distance, but it considers the covariance between dimensions/EEG channels, and can substantially improve decoding performance, in particular when used on EEG data with highly correlated EEG channels. The EEG data was first smoothed with a Gaussian smoothing kernel (SD=10 ms) along the time dimension and mean centered across channels prior to analysis. Using an 8-fold cross-validation approach, the trial-wise Mahalanobis distances between train trials and the left-out test-trials was computed at each time-point separately using the MATLAB function “pdist2” from the “Statistics and Machine Learning Toolbox”. The covariance matrix was computed at each time-point using all train trials with a shrinkage estimator using custom MATLAB code (Ledoit & Wolf, 2004). This procedure was repeated 50 times, each time randomly partitioning the data into 8 folds. Same-location distances were subsequently subtracted from the average of all different-location distances, such that a positive “distance difference” represents a shorter distance between trials that had the same missing square locations than between trials that had different locations, similar as was done previously for orientation decoding (Wolff et al., 2015), suggesting that the electrophysiological signal contains location-specific information. We ran this analysis separately for the three conditions of interest (SOA 0, SOA 30, and SOA 45), and for completeness repeated the same analyses including all 59 EEG channels.

For visualization purposes, we complemented the temporal decoding analysis with a searchlight decoding analysis where the decoding was repeated iteratively for a subset of electrodes across all 59 electrodes. For each iteration, the current as well as the closest two neighboring electrodes were included (similar to van Ede et al., 2019). This was done separately for each previously identified processing stage (90 to 140 ms, 140 to 300 ms, 300 to 500 ms, 500 to 1100 ms). Within each time-window, information was pooled across time to take advantage of the fact that information is also contained in the temporal profile of the data, for which we used a previously used approach (Wolff et al., 2020). That is, for a given time-window, the data of the three channels in question was first downsampled to 102.4 Hz (by taking the average of 10 consecutive time-points) and then added as additional dimensions to the decoding analysis. For example, for the early time-window (90 to 140 ms), there were 5 time-points per channel, which resulted in 15 dimensions for the subsequent decoding analysis (3 channels by 5 time-points), which was otherwise the same as described as above. The MATLAB extension fieldtrip (Oostenveld et al., 2010) was used to visualize the decoding topographies.

### Cross-temporal decoding

We were interested in the neural dynamics of the processing of the missing square. Instead of training and testing the classifier on the same time-points as described above, in cross-temporal decoding analyses, the classifier is trained and tested on all possible time-point combinations, resulting in a full cross-temporal decoding matrix (King & Dehaene, 2014). The cross-temporal dynamics were tested for significance using the same method as in Myers and colleagues (2015): The decodability at each cross-temporal time-point *tx,y* to the corresponding time-points on the diagonal (*tx,x* and *ty,y*). A significant difference in both was taken as evidence for dynamic coding. A cluster-based permutation test was used to correct for multiple comparisons.

Finally and most importantly, we wanted to test to what extent the processing of the missing square location is temporally shifted when the necessary visual information is not presented simultaneously, as in the SOA 0 condition, but spread across two discrete displays, as in the SOA 30 and SOA 45 conditions. To do so, we used the same decoding approach as described above, but instead of using cross-validation with separate train and test folds of the same SOA condition, we trained the classifier on the SOA 0 condition, and tested it separately on both the SOA 30 and SOA 45 conditions. As we did not know to what extent the visual processing of the missing square is temporally shifted between the SOA 0 condition and the two integration conditions (for example, it could be time-locked to the first or the second display, or somewhere in-between), we obtained the full cross-temporal decoding matrix as described above.

### Quantifying the temporal shift

We wanted to quantify the possible temporal shift obtained from the cross-temporal and cross-condition decoding matrices, where the classifier was trained on the no-integration condition (SOA 0) and tested on each temporal integration (SOA 30 and 45, see above). To quantify this shift, we developed an analysis that assumes that a dynamic, cross-temporal decoding profile is symmetric around the true temporal offset. First, the dynamics of the decoding matrices were normalized at each time-point by z-scoring the decoding values across both the x and y axes of the matrices and then averaging them. Then, a centered 50 ms sliding window approach was used to determine the temporal shift. At each time-point along the diagonal (from 90 ms, the onset of reliable decoding in the SOA 0 condition, to 500 ms), the average of all decoding values in the upper triangle were subtracted from the average of the lower triangle. This sliding window was moved up and down the training and testing axes of the decoding matrix. Zero difference at a specific offset indicates that the cross-temporal decoding window is symmetrical at this offset, whereas a positive or negative difference provides evidence for a temporal shift in the corresponding direction.

The “shift score” was computed using the following formula:

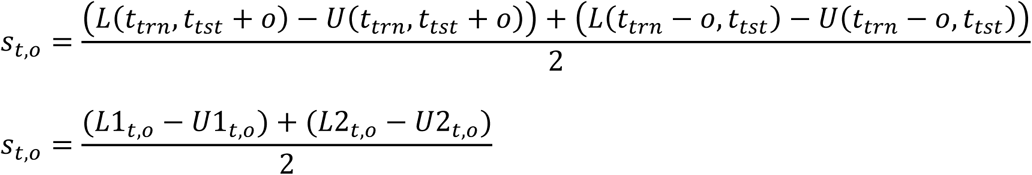

where *ttrn* and *ttst* are the time-points along the training and testing axis, respectively, both relative to the onset of the first display (*ttrn = ttst*), *U* and *L* are the means of the upper and lower triangle of the 50ms sliding window, respectively, *o* is the temporal offset of the sliding window (in ms) relative to *t,* and *s* is the shift score.

### Identifying distinct stages of location processing

The time-course of decoding and the cross-temporal decoding matrix of the no-integration condition were compared to the ERP, to identify different stages in processing of the location of the missing element. Stages were characterized on the basis of the correspondence observed between ERP component amplitude, decoding accuracy, and (switches between) periods of dynamic and stable coding. The identification of these stages was not necessarily meant to strictly delineate separable or independent processes, but to provide a handle on different moments within the perceptual process.

### SOA 0 to display 1 location generalization

In light of the results, it could be argued that display 1 alone in the temporal integration conditions provides partial information of where the missing element is going to be (i.e., all empty locations), before display 2 is even presented. This could in turn be picked up by the decoder as an earlier than expected signal of the missing element, which could (at least in part) occur before display 2 is even processed and temporally integrated with display 1 by the brain. To this end, we attempted to test if, and to what extent, the signal of the missing element location in the SOA 0 condition generalizes with empty locations in display 1 that are not the missing element in the integration conditions.

This would mean that the signal of the missing element location in the SOA 0 should be similar with the signal of trials from an integration condition (SOA 30 and 45) when the same location is empty at display 1, but is occupied by display 2 (meaning that it is not actually the missing element location in those trials), compared to trials where it is the other way around (location occupied at display 1, but empty at display 2; see Supplemental Fig. 4A for a schematic). Thus, if the empty locations in display 1 leads to a prediction of where the missing element could be that results in a measurable signal, which generalizes with signal of the SOA 0 condition, then the trials this applies to (display 1 empty) should be more similar (shorter Mahalanobis distance) to the corresponding SOA 0 trials than the trials this does not apply to (display 1 occupied and display 2 empty).

This is exactly what we tested. Specifically, from the SOA 0 condition we first computed the covariance matrix using all trials before the signal was averaged over trials based on the missing element locations (just like before). Next, the distance difference between display 1 empty and display 2 empty of a specific temporal integration condition was computed for all missing element locations. For example, say for the missing element location 5 in the SOA 0 condition, the signal of trials of a given integration condition were averaged based on whether location 5 was empty in display 1 *or* in display 2 (but never both). The distance to the average signal of each group of trials was then computed between the average signal over all missing element location 5 trials in the SOA 0 and the two trial-types of a given integration condition. Then, the distance difference was computed, where the distance to the display 1 empty trials was subtracted from the distance to the display 2 empty trials. The same was repeated for all 16 missing element locations, each time averaging the trials of the integration condition based on whether the corresponding location was empty at display 1 or 2, before the average distance difference (display 1 empty - display 2 empty) was obtained. This was done for each time point, where the signal of the SOA 0 condition was time-locked to the onset of the only display, and the signal of the integration conditions was time-locked to the onset of display 1. This procedure was done separately for the SOA 30 and the SOA 45 conditions.

### Significance testing

We used non-parametric permutation tests for all statistical analyses. In subject analyses, for each subject, the sign of the decoding or the deviation of the “shift-score” from the temporal offset in question was randomly flipped at each time-point 10,000 times. The resulting null-distribution was used to derive the *p*-value. Cluster-based permutation tests were used to correct for multiple comparisons across the time-dimension(s) using 10,000 permutations, with a cluster-forming threshold of *p* < 0.05. Statistical significance was set at *p* < 0.05. All tests were two-sided. Additionally, Bayes factors were obtained for the average “shift-score” deviations at each processing stage, as well as for each time-point and temporal offset (the latter only for visualization). For this we performed Bayesian *t*-tests with the default Cauchy prior scale of 0.707 (Rouder et al., 2009).

Decoding topographies were tested for significance using the cluster-based permutation for channels as implemented by the MATLAB extension fieldtrip (Oostenveld et al., 2010), where, instead of testing for significant decoding clusters over time (as described above), the topographies were tested for significant decoding clusters across space. The neighbors for each channel were first defined using the triangulation method of the fieldtrip function “ft_prepare_neighbours”. Then electrode clusters with significant decoding were obtained for each time-window of interest using the Monte Carlo method with 10,000 permutations, a cluster-forming threshold of *p* < 0.05, and a minimum of two neighboring channels for a sample to be included in the clustering.

Group-level analyses were used to estimate the temporal shift at each specific processing stage (detailed below) and SOA condition. The mean temporal shift values were sampled with replacement from each time-window of interest and the shift was computed 5,000 times. The resulting distribution was used to compute the 95% confidence interval.

Differences between SOAs and processing stages were also tested on the group level by randomly flipping the condition labels of each subject with 0.5 probability, before computing the mean difference 10,000 times. The null distribution was used to calculate the proportion of permutations more extreme than the actual group-level difference. Statistical significance was set at *p* < 0.05. Tests between SOA conditions were one-sided, testing specifically if the temporal shift in the SOA 45 condition was larger than in the SOA 30 condition. These tests were one-sided, as greater SOA would never be expected to reduce temporal shift, and vice versa. The *p*-values of all possible pairwise differences between processing stages were Bonferroni-corrected for multiple comparisons and were two-sided.

## Results

### Behavioural

Figure 1b shows a clear negative effect of increasing SOA on the behavioural accuracy of locating the missing square (*p* < 0.001 of all pairwise comparisons, two-sided, Bonferroni-corrected). This suggests that the two displays are less likely to be temporally integrated at longer SOAs, leading to worse performance, which replicates previous findings (Akyürek et al., 2010; Di Lollo, 1977).

### Distinct stages of location processing

We used the SOA 0 condition as a “template” to define the time-windows of individual processing stages, and to test the neural dynamics of location processing. The decoding time- course showed a significant cluster from 91 ms to 1,100 ms (end of epoch), relative to display onset (*p* < 0.001, cluster-corrected), providing evidence that the location of the missing square was decodable from the EEG shortly after stimulus presentation (Fig. 2a). The SOA 30 and 45 conditions showed similar but weaker decoding profiles (Supplemental Fig. 1), likely due to the weaker percept of the missing element and lower behavioral accuracy in these conditions.

**Figure 2.**
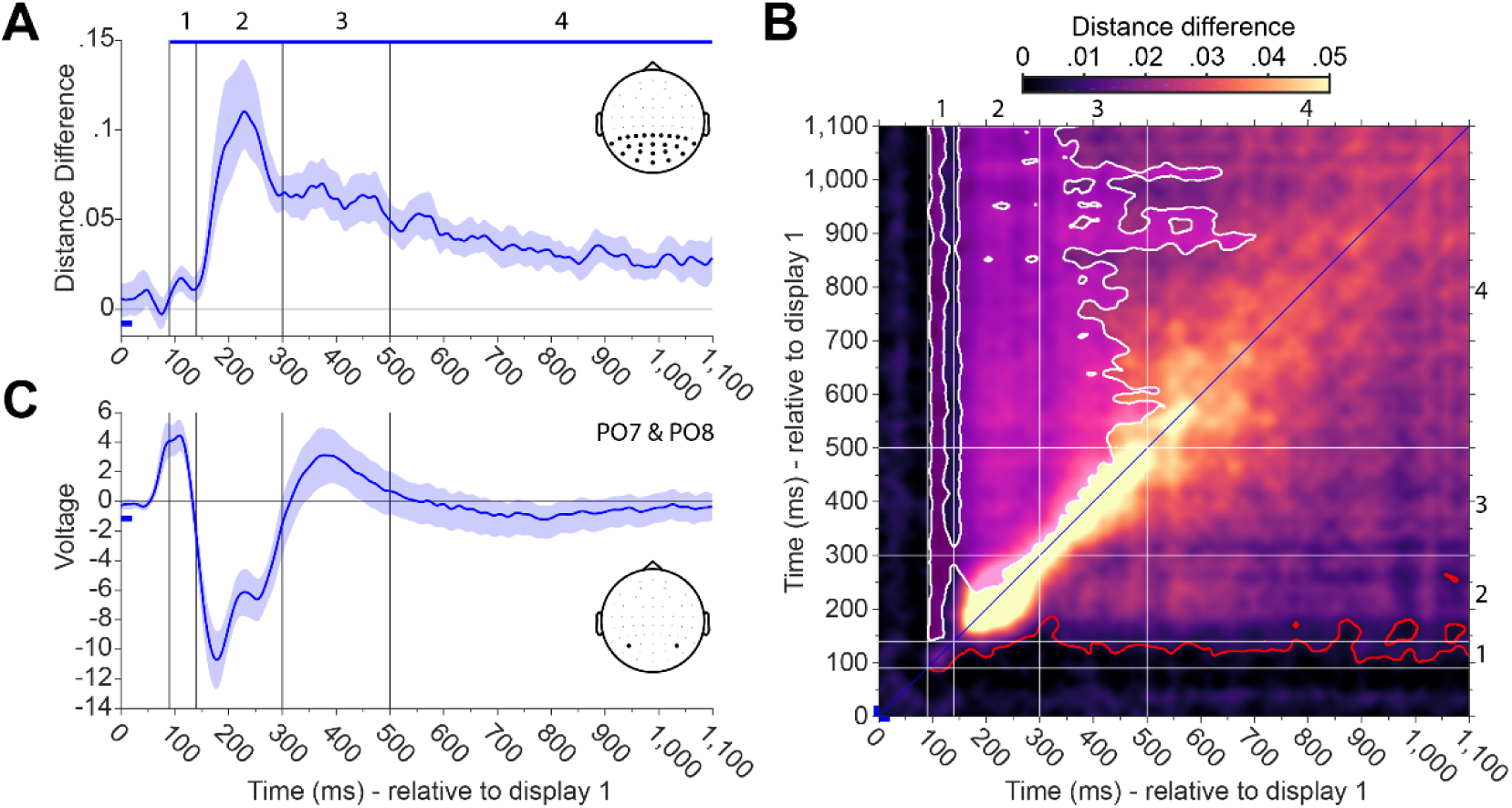
Identifying distinct processing stages (A) Decoding time-course of the location of the missing square in the SOA 0 condition using the posterior electrodes. Blue bar on top indicates significant decoding (p < 0.05, cluster- corrected). Blue bar on the bottom left indicates Display 1 presentation. Error-shading indicates the 95% C.I. of decoding accuracy. Black vertical lines and numbers indicate distinct processing stages (see text). (B) Cross-temporal decoding matrix of the SOA 0 condition. The white outline depicts time-points of significantly lower decoding than for both equivalent time- points along the diagonal, providing evidence for dynamic coding (p < 0.05, cluster-corrected). The red outline indicates time-points of significant decoding (p < 0.05, cluster-corrected). (C) Average evoked potential from the electrodes PO7 and PO8 with 95% C.I. (blue shading).

The cross-temporal decoding matrix of the SOA 0 condition showed that approximately 140 ms after stimulus onset training and testing trials cross-generalized significantly, while at the same time providing evidence for a highly dynamic code from decoding onset until approximately 500 ms after onset (decoding cluster: *p* < 0.001; dynamic coding cluster: *p* < 0.001, both cluster-corrected; Fig. 2b). Visual inspection of the 1-dimensional and 2- dimensional decoding time-courses, in conjunction with the significant clusters of decoding and dynamic coding, suggested four distinct processing stages, which corresponded well with the averaged evoked potentials (Fig. 2c).

1. An early location-specific response from approximately 90 to 140 ms, which did not cross-generalize with any subsequent time-points. The decoding time-course overlapped with the P1, which is known to be modulated by early attentional filtering and stimulus selection (Hillyard et al., 1998), and which originates in V2 (Woldorff et al., 1997).
2. High decodability from 140 to 300 ms, overlapping with the time-course of the N1 and N2, likely reflecting the attentional shift towards the location of the missing square (Patel & Azzam, 2005), and spatial grouping (Akyürek et al., 2010).
3. Highly dynamic location decoding from 300 to 500 ms, overlapping with the time-course of the P3, likely reflecting the consolidation of the missing square location in working memory, and processes related to response selection (Polich, 2007; Verleger et al., 2005).
4. Stable and relatively low decoding from 500 to 1,100 ms (end of epoch), likely reflecting the continuous maintenance of the missing square location in working memory, possibly related to the CDA (Akyürek et al., 2017; Vogel & Machizawa, 2004).

Since we were interested in the temporal shift of the dynamic brain states between no-integration and integration, we focused on the dynamic time-window up to 500 ms after stimulus presentation for all subsequent analyses, excluding the stable maintenance period.

Searchlight decoding analyses over electrode-clusters of the SOA 0 condition for each processing stage showed that decoding was generally most robust in posterior electrodes (Supplemental Fig. 2), this, in conjunction with lower decoding when using all electrodes for decoding (Supplemental Fig. 1B), corroborates our a-priori choice to only use the posterior electrodes for the main analyses.

### Exemplifying “shift score” computation on SOA 0

We quantified the mid-point of the cross-temporal decoding matrix relative to the diagonal by computing the shift score. Here we demonstrate this analysis on the cross-temporal decoding matrix of the SOA 0 condition, where we do not expect a systematic deviation from diagonal, since both training and testing come from the same SOA 0 condition. The method is described above (“Quantifying the temporal shift”) and illustrated in Figure 3. A significance test on the shift-score matrix reveals one negative and one positive cluster of shift-score values (*p* < 0.001, cluster-corrected; Fig. 3b). Neither of the clusters cross the 0 line, demonstrating that while there is no evidence for a temporal shift when training and testing at the same time- points, the shift-score analysis is sensitive enough to pick up the asymmetry of the decoding matrix when separating training and testing by as little as ∼5 ms. The 0 ms offset was explicitly tested for significant deviations of the shift-scores. As expected, no significant clusters were detected (*p* > 0.2, cluster-corrected; Fig. 3c).

**Figure 3.**
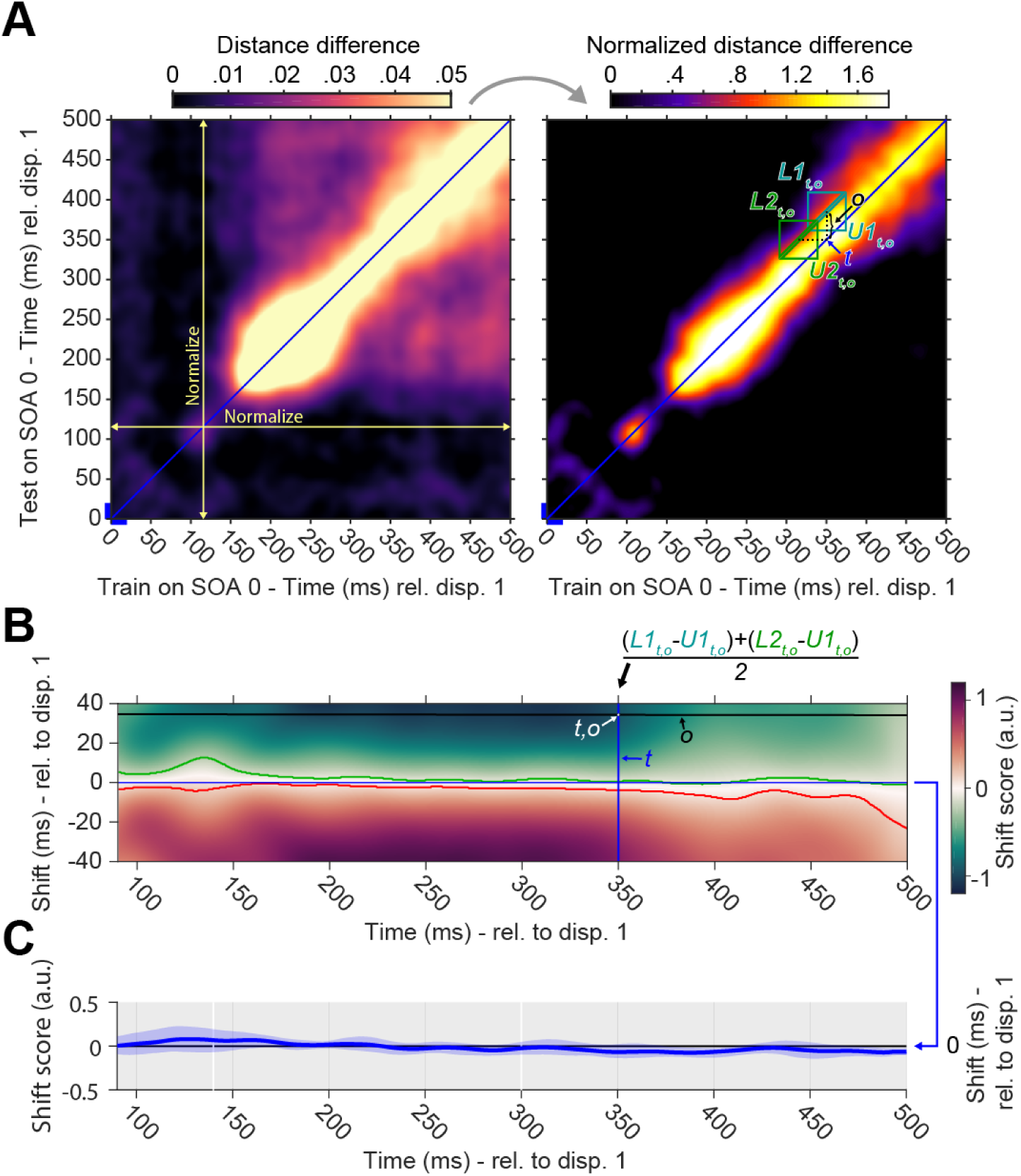
Demonstration of “shift score” computation (A) Cross-temporal decoding matrix of the SOA 0 condition (left). Yellow lines visualize normalization across cross-temporal time-dimension. Normalized decoding matrix by z-scoring decoding values along the cross-temporal time-dimensions (right). Computation of shift score at a specific time (t) and temporal offset (o) is illustrated. (B) Shift scores with temporal offsets from -40 to 40 ms. Negative values suggest higher decodability at shorter offsets, positive values suggest higher decodability at longer offsets. Green and red outlines depict significantly negative and positive shift-score clusters (p < 0.05), respectively. (C) Shift score at 0 ms temporal offset with 95 % C.I. No evidence of significant deviation was found (p > 0.2, cluster-corrected), suggesting that the neural dynamics are centered at 0 (as expected).

### Temporally shifted neural dynamics of location processing during temporal integration

The neural dynamics of missing square location processing in the SOA 0 condition cross-generalized significantly with both temporal integration conditions (*p* < 0.001 of both cross-temporal decoding clusters; Fig. 4a & b), with similar neural dynamics across the whole time-window as SOA 0 (see Fig. 3a).. However, a temporal shift seemed to be present in both temporal integration conditions, as the decoding dynamics were not symmetrical along the diagonal (training at testing at the same time-points, relative to Display 1 onset).

**Figure 4.**
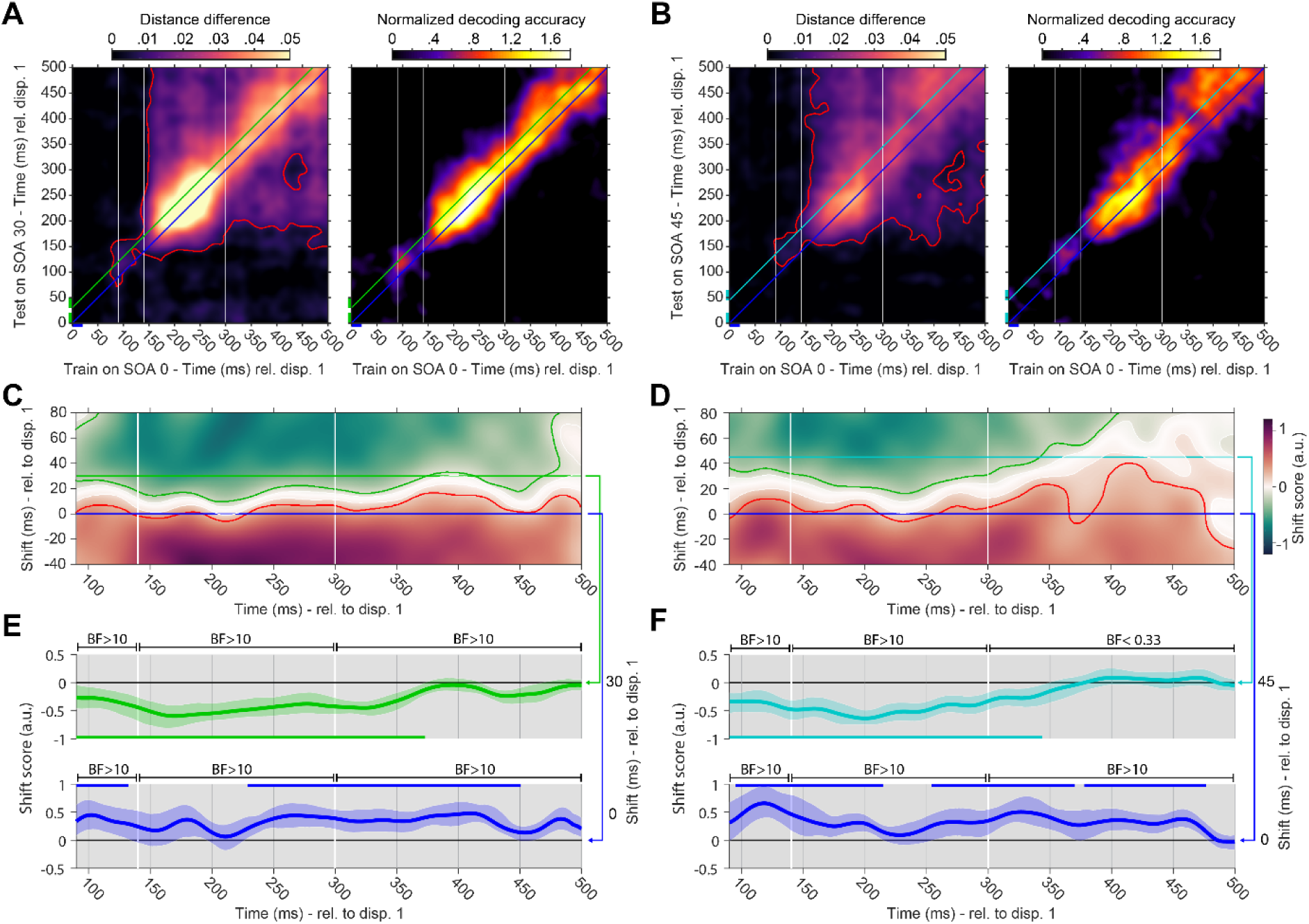
Accelerated location processing during temporal integration (A) Cross-temporal generalization matrix between SOA 0 and SOA 30. White lines indicate previously defined processing stages. Left: The red outline indicates time-points of significant cross-generalization (p < 0.05, cluster-corrected). Right: Normalized generalization matrix. (B) Cross-temporal generalization matrix between SOA 0 and SOA 45. (C) and (D) Shift scores of normalized cross-temporal generalizations between SOA 0 and SOA 30, and SOA 0 and SOA 45, respectively. Green and red outlines depict significantly negative and positive shift-score clusters (p < 0.05), respectively. White outlines and shading show shift scores with strong evidence against a positive or negative deviation (i.e., evidence for no deviation), representing the estimated temporal shift of processing (Bayes factor < 0.333). (E) Shift scores of SOA 30 at 30 ms (green) and 0 ms (blue) temporal offset. Bars in the corresponding colors show significant shift scores (p < 0.05, cluster-corrected). Error-shading indicate 95 % C.I. of the mean. Approximate Bayes factors (BF) are shown for the average shift-scores of each processing stage (F) Shift scores of SOA 45 at 45 ms (cyan) and 0 ms (blue) temporal offset. Same convention as E.

Two clusters of significant shift-scores were present in the SOA 30 condition, one negative and one positive (both *p* < 0.001, cluster-corrected; Fig. 4c). For the most part, the point of decoding symmetry (shift-score = 0) was wedged between the onset of Display 1 (0 ms) and Display 2 (30 ms), constituting evidence for perceptual acceleration during temporal integration. A positive and negative shift-score cluster were also present in the shift score matrix of the SOA 45 condition (both *p* < 0.001, cluster-corrected; Fig. 4d). Like SOA 30, the point of symmetry was largely between the onsets of the first and second display, though the acceleration during integration seemed to diminish towards the end of the dynamic time- window.

We explicitly tested if there was evidence for a temporal shift at the temporal onsets of the first (0 ms) and the second display (30/45 ms) in both temporal integration conditions. In the SOA 30 condition, this revealed a significant negative shift-score cluster at a temporal offset of Display 2 onset/30 ms (*p* < 0.001, cluster-corrected; Fig. 4e, top), providing evidence for a temporal offset of the neural dynamics that is less than the SOA between displays (30 ms). Two positive shift-score clusters were present at 0 ms temporal offset (*p* = 0.043 and *p* < 0.001, cluster-corrected; Fig. 4e, bottom), indicating that location processing is delayed during temporal integration relative to Display 1 onset. The SOA 45 condition showed a similar pattern of results. One negative shift-score cluster was present at a temporal offset of Display 2/45 ms (*p* < 0.001, cluster-corrected; Fig. 4f, top), as well as three positive clusters at 0 ms temporal offset (*p* < 0.001, *p* < 0.001, and *p* = 0.004, cluster-corrected; Fig. 4f, bottom).

Bayesian analyses performed on the average shift-scores over the temporal windows of each previously defined processing stage showed strong evidence for positive shift-scores in the SOA 30 condition relative to a 0 ms temporal offset (90 to 140 ms, *BF* = 19.139; 140 to 300 ms, *BF* = 55.147; 300 to 500 ms, *BF* > 10000), and strong evidence for negative shift- scores relative to a 30 ms temporal offset (90 to 140 ms, *BF* = 27.586; 140 to 300 ms, *BF* > 1000; 300 to 500 ms, *BF* > 1000). The results for the SOA 45 condition showed strong evidence for positive shift-scores relative to a 0 ms offset (90 to 140 ms, *BF* = 63.831; 140 to 300 ms, *BF* > 1000; 300 to 500 ms, *BF* > 1000). While there was also strong evidence for negative shift scores relative to a 45 ms offset for the first two stages (90 to 140 ms, *BF* = 170.117; 140 to 300 ms, *BF* > 1000), there was evidence *against* a shift-score deviation for the last stage of interest (300 to 500 ms, *BF* = 0.278).

In sum, when Display 2 was delayed, processing of the missing element location was also delayed, compared to the SOA 0 condition (Fig. 4e & f, bottom). However, it was not delayed as much as would be expected from the SOA (i.e., 30 or 45 ms) between displays (Fig. 4e & f, top). That means that the processing of the missing element was actually accelerated in these integration conditions, in particular during the early processing stages.

### Distinct temporal shifts of location processing stages during temporal integration

Average temporal shifts at each previously defined processing stage are shown in Table 1 and Figure 5. Apart from one exception, all confidence intervals of the temporal shifts fell in-between the onsets of Display 1 and 2. Group analyses showed that while the temporal shift was not significantly higher in the SOA 45 condition than the SOA 30 condition in the early processing stage (90 to 140 ms, *p* = 0.099), its average shift was higher in both the middle (140 to 300 ms, *p* = 0.021) and late processing stage (300 to 500 ms, *p* < 0.001).

**Figure 5.**
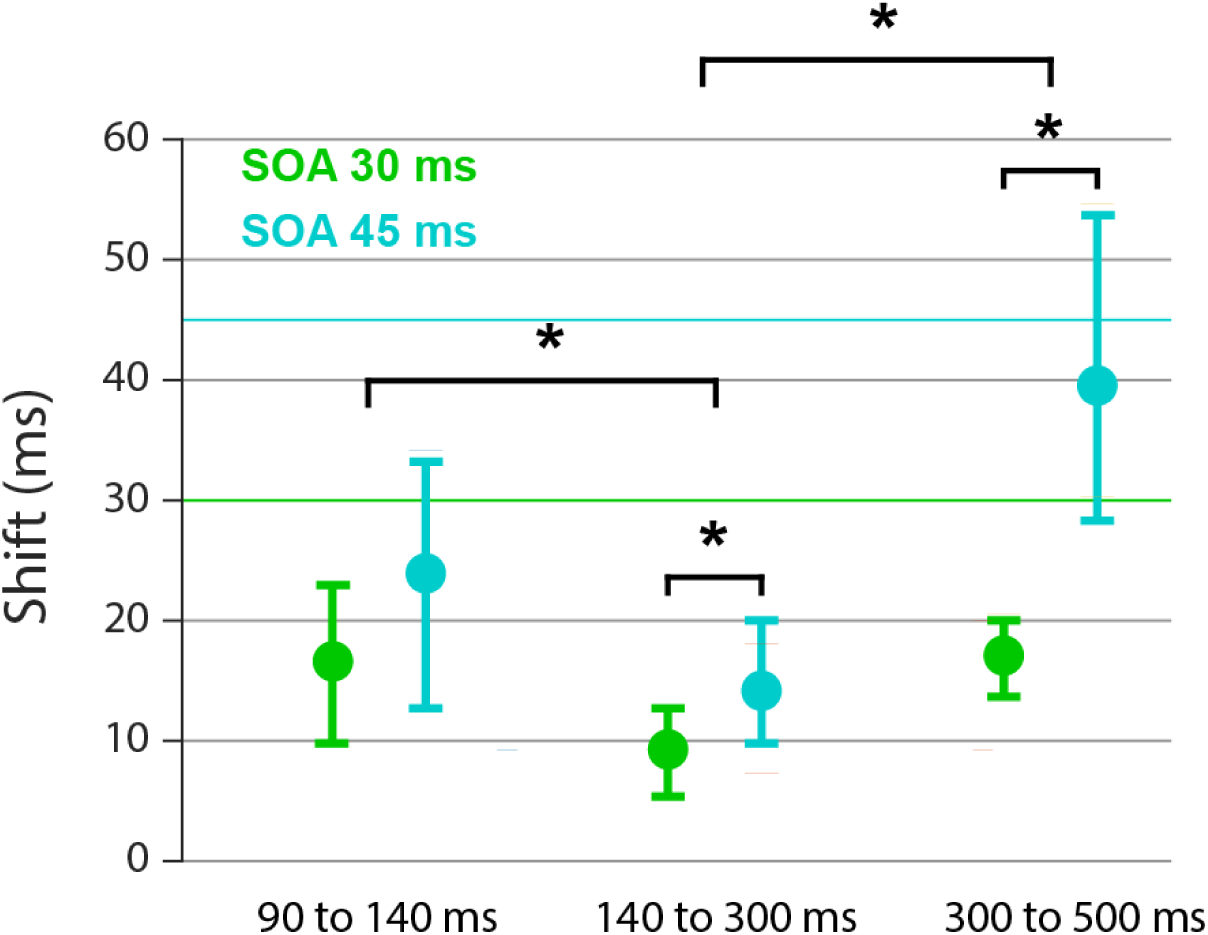
Temporal shifts of location processing dynamics Error-bars are 95% C.I.s. Asterisks denote significant differences between SOAs and processing stages (*p* < 0.05, Bonferroni-corrected for comparisons between stages).

**Table 1.** Temporal shifts of location processing dynamics as a function of temporal integration condition (SOA 30/45), for each processing stage. 95% Confidence intervals are shown in brackets.

| Time<br>Condition | 90 to 140 ms | 140 to 300 ms | 300 to 500 ms |
| --- | --- | --- | --- |
| SOA 30 | 17 ms<br>(10-23 ms) | 10 ms<br>(6-13 ms) | 18 ms<br>(14-21 ms) |
| SOA 45 | 24 ms<br>(13-33 ms) | 15 ms<br>(10-20 ms) | 40 ms<br>(28-55 ms) |

Pairwise comparisons between processing stages (averaged across both conditions) revealed that the temporal shift at the middle processing stage was significantly lower than at both the early and late processing stages (*p* = 0.004 and *p* = 0.001, respectively, Bonferroni-corrected), while the temporal shifts at the early and late stage did not differ significantly (*p* > 0.5, Bonferroni-corrected).

Results obtained when using all EEG channels for the previous decoding procedures were qualitatively similar, though the difference between temporal shifts of the SOA conditions at the middle processing stage was not significant (Supplemental Fig. 3).

### No evidence that partial display 1 information generalizes with SOA 0 decoding

It could be argued that the previously reported results may not necessarily be due to accelerated perception during visual temporal integration, but rather because the presentation of the first display alone already provides partial information about where the missing element is going to be. If so, the signal associated with empty locations at display 1 in the integration conditions (SOA 30 and 45) should generalize with the signal of the missing element location in the SOA 0 condition (for a more detailed explanation, see methods and Supplemental Fig. 4A). However, we found no evidence that this was the case for either the SOA 30 or the SOA 45 conditions (Supplemental Fig. 4B).

## Discussion

We decoded the location of the missing element from EEG recorded in the temporal integration task that bears its name. This location is only apparent when the two successive stimulus displays in the MET are temporally integrated into a single perceptual representation, thus providing a highly time-sensitive fingerprint that can be tracked in the brain. We observed that perception is accelerated during temporal integration: The location of the missing element could be decoded sooner than would be expected on the basis of the actual delay between the stimulus displays. Compared to simultaneous presentation of the displays, the missing element was delayed by only 15 ms on average at an SOA of 30 ms (corresponding to an acceleration of 15 ms also). At an SOA of 45 ms, the missing element was delayed by only 26 ms on average (19 ms acceleration). These results constitute evidence that the perception of the first display facilitated that of the second, thereby possibly aiding in their joint integration. This evidence of perceptual acceleration during temporal integration places our fast-paced task within a larger family of spatial/temporal cueing (Coull & Nobre, 1998; Posner, 1980), rapid resumption (Lleras et al., 2005; Spaak et al., 2016), foreperiod (Luce, 1991), and rhythmic tasks (Jones et al., 2002; Mathewson et al., 2010), in which advance information aids the observers in perceiving and acting more quickly.

Importantly, we also observed that acceleration varied depending on the precise moment in the perceptual processing pathway. Combined across SOAs, processing was accelerated by 17 ms in the early stage, by 25 ms in the middle stage, and by 9 ms in the late stage. Thus, perceptual acceleration was neither linear, nor constant, but variable to the extent that early gains were at least partially lost again later. The observed non-uniformity might be conceptualized as a queuing effect that may happen when information is passed from one stage to another, such that rapid completion of processing in one stage may be followed by increased waiting time for the next stage. In other words, as information flows from one processing stage to the next, it can wax as well as wane.

### The locus of perceptual acceleration

In our experimental task, processing the second display was facilitated only by the onset of the first, less than 50 ms prior, which is an interval that has been considered too short to afford proper temporal preparation (Los & Schut, 2008). Indeed, at this temporal offset, the acceleration effect would seem to be driven purely by low-level factors within local neuronal assemblies in early visual cortex, such as synaptic facilitation (Markram et al., 1997; Zucker & Regehr, 2002). From this perspective, it is all the more surprising that the acceleration effect on the resultant representation was not straightforward (i.e., constant or linear). Such variable acceleration might have been more expected to result from higher-level factors (e.g., stimulus familiarity), which might arrive too late to interact with all processing stages, rather than the present manipulation.

The waxing and waning of acceleration that we observed during temporal integration is at odds with simplistic models of perceptual processing. In particular models with few parameters, such as (drift diffusion) models of temporal preparation, which allow only for alterations of the rate at which sensory evidence accumulates increase, and of the onset of this process (e.g., Seibold, Bausenhart, et al., 2011; Vangkilde et al., 2012; see also Jepma et al., 2012), will be unable to account for the variable pattern we presently observed. Allowing some interaction between parameters would seem to be necessary.

More generally, this limitation also applies to traditional additive factors logic (Donders, 1868; Linden, 2007; Sternberg, 1969, 2001). In this framework, the chain of processing is divided into distinct, independent stages or modules, each of which is conceptualized as an independent computation, transforming input to output. Acceleration may act on each stage, reducing the time needed to complete that transformation, without affecting any other processing stage. The relationship between different stages is also summative, so that subsequent computations in the perceptual and cognitive chain will commence sooner, until eventually the behavioral response is also quickened. Although attractive for its elegant simplicity, this framework cannot accommodate the variable effects that we found.

It may be noted here that our study is not the first to question the idea of independence between different processing stages. Under- and over-additive effects have previously emerged in dual-task situations, where the processing of multiple (conflicting) stimuli is required simultaneously (Ridderinkhof et al., 1995), and where multiple subtasks need to be chained (e.g., Sackur & Dehaene, 2009), but also in single-task foreperiod studies (e.g., Los & Schut, 2008). These findings also suggest that concurrent modulation of multiple stages can occur (Marti & Dehaene, 2017; see also Caplan, 2007 for an extended discussion). Recent stage-based models have incorporated this by allowing temporal variation within each stage, and by using neurophysiological markers (“bumps”) to define the stages as well as their number (Anderson et al., 2016; van Maanen et al., 2021; Vidaurre et al., 2019, 2021). The loss of simplicity in these models may detract to some degree from the idea of a serial processing chain, and make it harder to pinpoint modulatory factors therein. Overall, however, these more data-driven stage-based models may currently hold the most promise in explaining perceptual acceleration as well.

As a caveat, it must be noted that the stages we identified in the present data were not the result of systematic analyses of the EEG, because it was not our purpose to attempt such identification. We therefore do not want to overstate the stage-specific nature of the observed acceleration effects. In our study, the processing stages have an arbitrary element. Their main purpose was to provide a handle on different moments in what might as well be considered a continuous processing stream (cf. Fig 4c & d). Importantly, the conclusion that the representation of the missing element location exhibited varying degrees of acceleration across the interval under consideration is largely unaffected by whether stage boundaries might be drawn differently within.

### Temporal integration

The MET is traditionally used to measure visual temporal integration, and we indeed observed the associated pattern of results, in which increasing SOA resulted in fewer correct reports of the missing element location. The electrophysiological results provided an intriguing view on how temporal integration may be facilitated in the brain. To reiterate, the missing element location emerged sooner than would be expected on the basis of the SOA between the displays. This suggests that the processing of the first and second stimulus gets temporally ‘compressed’ in the brain. This compression is not predicted by models of temporal integration, which generally assume that integration is mostly a consequence of temporal overlap or coincidence of the processing of the second display with lingering representations of the first display, referred to as visible persistence, rather than any perceptual acceleration (Dixon & Di Lollo, 1994; Hogben & Lollo, 1974; Loftus & Irwin, 1998; Sperling, 1960). Nevertheless, the current results suggest that perceptual acceleration may cause increased temporal overlap between stimulus representations when processing rapid successive stimuli, which would increase the chance that they are temporally integrated. Furthermore, the early arrival of the (features of the) trailing stimulus may facilitate the perception of simultaneity and thereby enhance temporal integration as well. Whether this perceived simultaneity is an independent factor might be a topic for further study.

## Conclusion

In the MET, we found that perception of the first display accelerates the processing of the second, which may contribute to the process of temporal integration. Temporal generalization analyses of the EEG patterns associated with the representation of the missing element showed that this acceleration is neither linear nor constant, and varies from moment to moment in the perceptual processing chain. Although the results are still compatible with the general idea of a fixed chain of perceptual processing stages, they do suggest that attempts to isolate and time processing stage-specific effects may mischaracterize the actual flow of events.

## Data and Code availability

All data are openly available on <u>osf.io/kftjw</u>. The custom Matlab code that reproduces all results in this manuscript is available on <u>github.com/mijowolff/Accelerating-perception-during-temporal-integration</u>.

## Author Contributions

Michael J. Wolff: Conceptualization, Methodology, Software, Validation, Formal analysis, Investigation, Data curation, Writing—original draft, Writing—review & editing, Visualization. Elkan G. Akyürek: Conceptualization, Methodology, Writing—original draft, Writing—review & editing, Supervision, Project administration, Resources

---

**Supplemental Figure 1.**
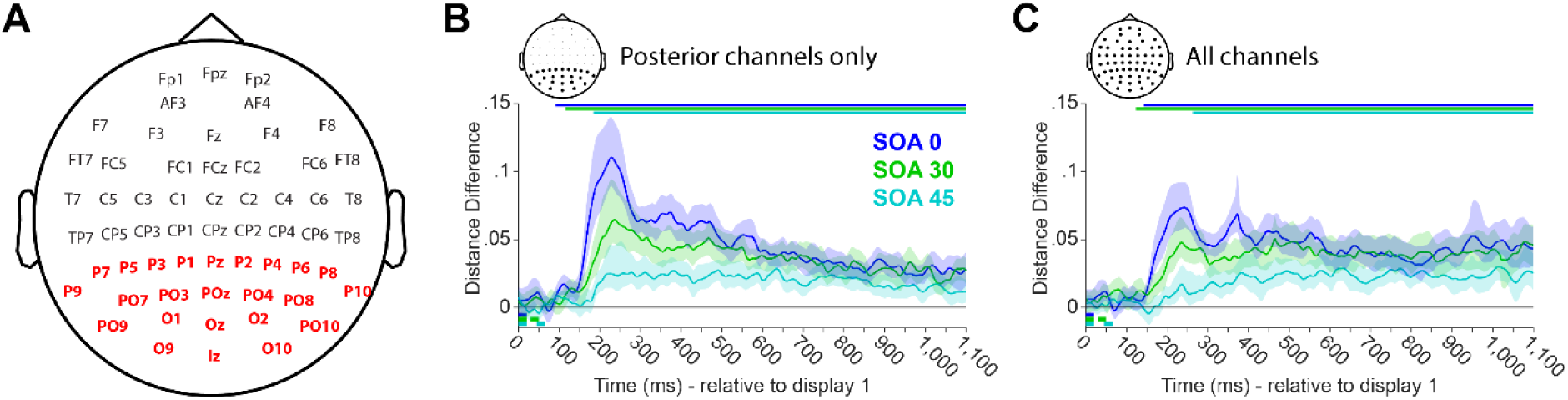
Channel layout and within-condition decoding for all conditions. **(A)** Channel layout and names of recorded EEG channels. Channels used for the main analyses are highlighted in red. **(B)** Decoding time-course of the location of the missing square in the SOA 0 (blue), 30 (green), and 45 (cyan) conditions, when training and testing within condition and time-point and using the posterior electrodes. Colored bars on top indicate significant decoding (*p* < 0.05, cluster-corrected) of the corresponding condition. Small colored bars on the bottom left indicate the display timings of the corresponding conditions. Error-shading indicates the 95% C.I. of decoding accuracy. **(C)** Same as (A) but using all electrodes for decoding.

**Supplemental Figure 2.**
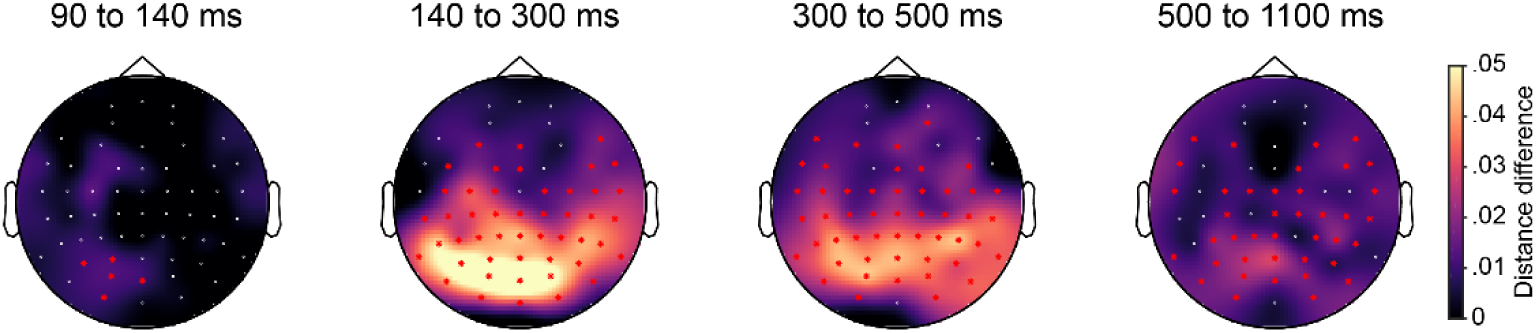
Decoding topographies of the SOA 0 condition. Decoding topographies of the searchlight analysis for each time-window. Significant electrode- clusters are highlighted with red asterisks (*p* < 0.05, cluster-corrected).

**Supplemental Figure 3.**
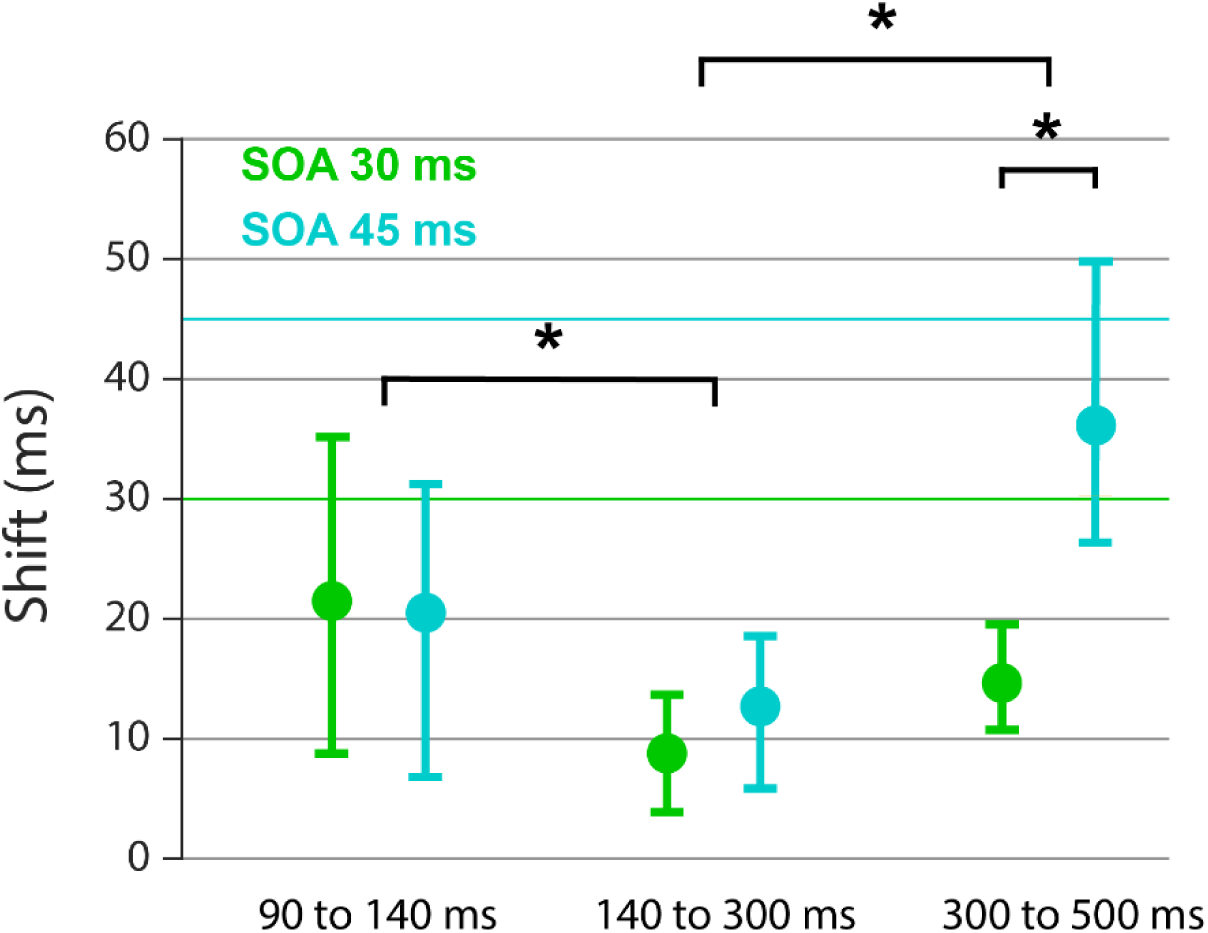
T**e**mporal **shifts of location processing dynamics when using all channels for decoding.** Same convention and analysis as in Figure 5, but using all EEG channels for the decoding analyses. Error-bars are 95% C.I.s. Asterisks denote significant differences. Pairwise average differences between processing stages (Bonferroni corrected for multiple comparisons): 90-140 ms versus 140-300 ms, *p* = 0.042; 140-300 ms versus 300-500 ms, *p* < 0.001; 90-140 ms versus 300-500 ms, *p* > 0.5. Differences between SOAs at each processing stage: 90-140 ms, *p* = 0.493; 140-300 ms, *p* = 0.096; 300-500 ms, *p* < 0.001.

**Supplemental Figure 4.**
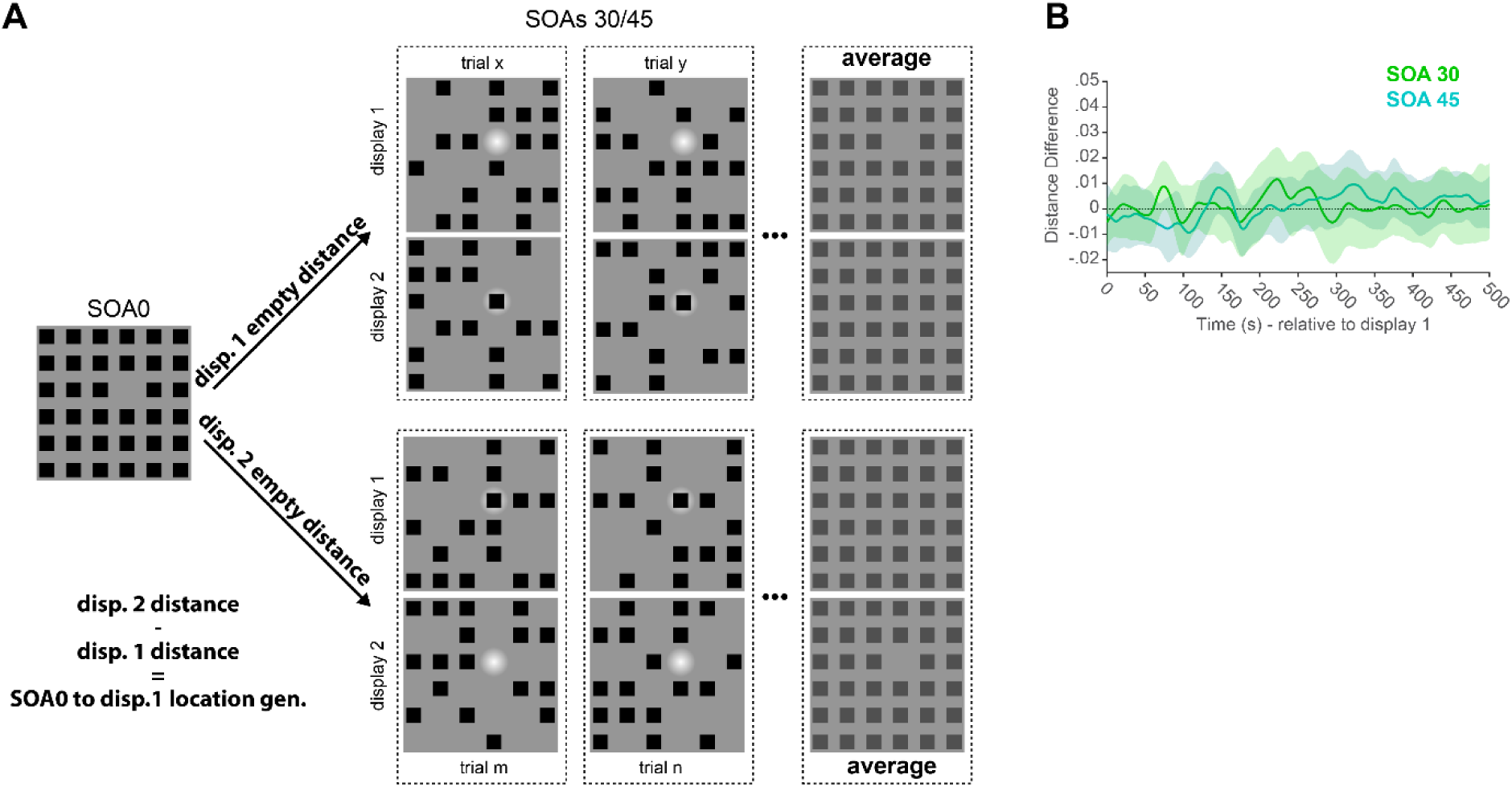
SOA 0 to display 1 location generalization**. (A)** Schematic of how the influence of partial information in display 1 on missing element generalization between conditions was tested. In short, for a given missing element location condition in the SOA 0 condition, trials of the integration conditions were grouped and averaged based on whether the corresponding location was empty in display 1 (top, trials “x” and “y”) or in display 2 (bottom, trials “m” and “n”). The distances to each group were computed. This was repeated for all missing element locations before all “display 1 empty distances” were subtracted from all “display 2 empty distances”, resulting in a distance difference that represents “SOA 0 to display 1 location generalization”, which is supposed to capture if, and to what extent, empty locations in display 1 in the integration conditions generalize to the missing element locations in the SOA 0 condition. **(B)** Results of the “SOA 0 to display 1 location generalization” procedure. No significant clusters in the generalization time-course of either integration condition (green: SOA 30, cyan SOA 45). Error-shading indicates the 95% C.I. of generalization accuracy.

